# A Galaxy-based training resource for single-cell RNA-seq quality control and analyses

**DOI:** 10.1101/724047

**Authors:** Graham J Etherington, Nicola Soranzo, Suhaib Mohammed, Wilfried Haerty, Robert P Davey, Federica Di Palma

## Abstract

**Background:** It is not a trivial step to move from single-cell RNA-seq (scRNA-seq) data production to data analysis. There is a lack of intuitive training materials and easy-to-use analysis tools, and researchers can find it difficult to master the basics of scRNA-seq quality control and analysis.

**Results:** We have developed a range of easy-to-use scripts, together with their corresponding Galaxy wrappers, that make scRNA-seq training and analysis accessible to researchers previously daunted by the prospect of scRNA-seq analysis. The simple command-line tools and the point-and-click nature of Galaxy makes it easy to assess, visualise, and quality control scRNA-seq data.

**Conclusion:** We have developed a suite of scRNA-seq tools that can be used for both training and more in-depth analyses.

## Background

The advent of RNA-seq has enabled a host of important discoveries in many biological areas such as gene expression, alternative splicing, comparative genomics, and gene annotation. Bulk RNA-seq, where a population of cells is used in every sample, usually provides copious amounts of RNA, but only measures the average expression level across that population. If different cell populations are included in a single sample, then information may be missed due to dominance of the transcription profile of one population against another. A decade ago, the development of single-cell RNA-seq (scRNA-seq) [1] made it possible to sequence the transcriptome of individual cells. This innovation opened the door to identification of novel cell types, uncovering regulatory pathways between genes, tracing the trajectories of distinct cell lineages, and pseudo-time reconstruction.

Typically, reads generated from cells in scRNA-seq experiments are mapped to a reference genome and then an expression matrix, calculated from the number of reads that are allocated to each gene or transcript, is produced. There are a number of tools available for this process (or variations of it) that are widely used within the community [2–4].

Along with the advantages of scRNA-seq come a number of technical challenges. scRNA-seq data is inherently ‘noisy’. Disruption or damage to the cell can result in the escape or degradation of nuclear DNA leaving predominantly cytoplasmic DNA in the cell. Further, inefficient RNA capture combined with amplification bias may distort gene expression profiles. This often results in ‘dropouts’, where genes are found to be at least moderately expressed in a few cells, but absent from most.

A large array of tools are now available to address quality control (QC) and expression analysis in scRNA-seq data (e.g. [5–11]).

The Scater package [12] provides the ability to QC scRNA-seq data, providing methods for visualisation, filtering, and expression analyses, as well as being compatible with further downstream analysis tools. Like many of the other tools cited above, Scater requires at least some experience of the Linux command line or even more advanced experience of the R programming language [13]. Also, due to both the large amount of sequencing data that scRNA-seq may produce and the high computational resources required by many of the tools, specialised infrastructure such as High-Performance Computing (HPC) is often required to analyse such experiments.

There is a lack of training resources for wet-lab scientists who want to carry out computational analyses of NGS data. Despite online resources for programming being quite plentiful along with a plethora of online help forums, resources focused on both training and analyses of biological data are few [14].

Galaxy is an open-source scientific workflow, data integration, and data analysis platform that aims to make life science research accessible to research scientists that do not have computer programming or systems administration experience [15]. Galaxy is available through well over one hundred public Galaxy servers, or can be easily integrated with existing HPC and cloud resources.

Over 30 scientific groups involved in Galaxy-related training contribute to the Galaxy Training Network (GTN) (https://galaxyproject.org/teach/gtn/). The GTN provides online training materials as well as coordinating Galaxy training events worldwide [16].

## Results

### Scater wrappers and workflows

Here we describe our Scater command-line wrappers, along with their integration into Galaxy and use on publicly-available Galaxy servers, for use as both an introduction to scRNA-seq data analyses and for more focussed downstream analyses.

Using Scater v1.10.1 [12], we defined the most common and intuitive tasks researchers new to the field of scRNA-seq might carry out in their analyses and created generic tools to accomplish these tasks. Typically, one would read in an expression matrix, calculate some metrics on the data, visualise the data and then filter out low quality cells or unexpressed genes. Then a further round of metrics would be calculated and visualised to assess the impact of the previous filtering steps. This visualise-filter-visualise iteration can be continued until a user is happy they have retained only high-quality data. The next step would be to look for confounding factors in the data, such as batch effect, by clustering the data and plotting it in relation to experimental meta-data, or any other non-biological variables that might have an effect on the final data. Such factors could include which plate each cell was generated on, the sequencing run, extraction date, lab technician, batch, replicate, etc.

Using methods from the Scater platform, along with other bespoke analysis and plotting methods, we have interpreted these tasks into a number of easy-to-use command line scripts, which requires only the most basic familiarity with the command line. Further, these scripts may be integrated into the Galaxy platform by using the Galaxy wrappers developed alongside the scripts. We also provide the inbuilt Scater data as a range of input files for users to input and test the methods below.

The basic workflow is as follows:

Step 1. The user inputs the data (a sample x gene read-count matrix), along with other metadata such as the experimental annotation and any control genes (often ERCC spike-ins or the names of the mitochondrial genes). This is then loaded into Scater and a number of quality control metrics are automatically calculated on the data. The output from this is a Loom file (http://loompy.org/), an HDF5-based format which is designed to efficiently store large omics datasets. These Loom files are then used as the input for all subsequent steps in the workflow.
Step 2. The data is visualised using a range of plots to show information about each cell. The distribution of reads in each cell, the number of genes expressed in each cell, and a scatter plot of the number of reads versus the number of expressed genes are all plotted. Finally, the percentage of mitochondrial genes in proportion to the total number of genes expressed is also plotted (Figure 1). These visualisations provide insight into poor quality cells that have either low read counts, low gene counts, or high mitochondrial gene expression.
Step 3. The data can now be filtered in two different ways. The user can decide to use information from the visualisations to, for example, remove cells that have low read count, or the user can use a PCA filtering method where cells calculated to be outliers are removed automatically. The input to this step (and in step 2 above) is the output from step 1. The output from this step is a new Loom file with the low-quality filtered-out cells removed.
Step 4. The filtered data can now be visualised (as in step 2 above) to assess the filtering process carried out in step 3. These two steps can be carried out iteratively, steadily increasing the parameters until the user is satisfied they have only the highest quality cells remaining (Figure 2).
Step 5. Once the user has a high-quality dataset, they can investigate any confounding factors in the data, such as batch effect. Any metric in the experiment annotation file may be plotted and variation in metrics or categories may be displayed by setting the size, colour or shape of the plotted points.

**Figure 1.**
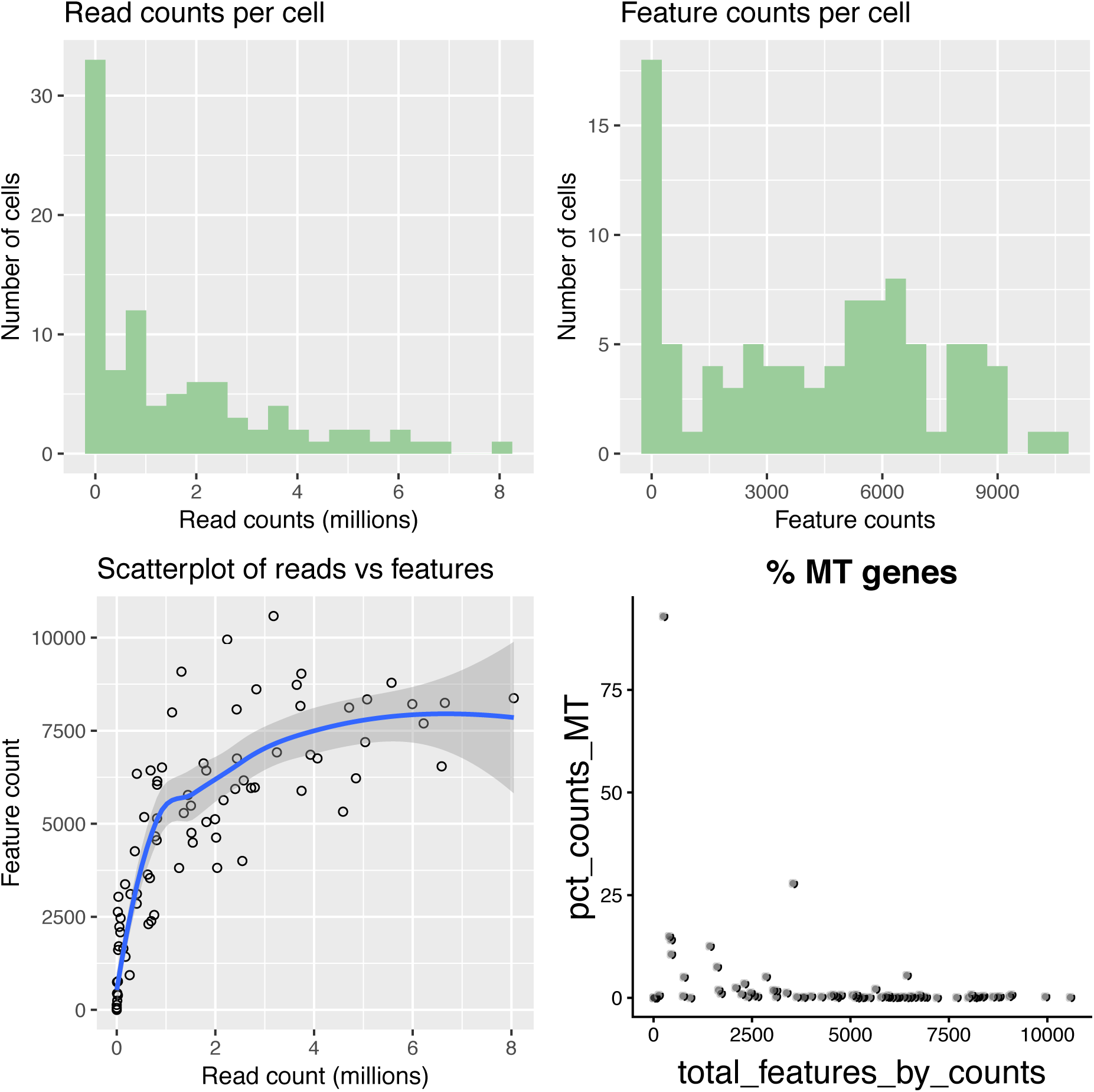
The plotting tools provide the ability to visualise the quality of each cell in the shape of histograms and scatter-plots. Here raw data, before being filtered is plotted. Typically, raw data will show a high number of cells with both low read counts (Read counts per cell) and low feature counts (Feature counts per cell). Low quality cells often have both low read and feature counts, depicted by the cluster of cells at the base of the x and y axes in ‘Scatterplot of reads vs features’. Lastly, the ‘% MT genes’ plot shows the proportion of reads mapping to mitochondrial genes. We can see that at least one cell has more than 75% of its reads mapped to mitochondrial genes, suggesting the cell was degraded.

**Figure 2.**
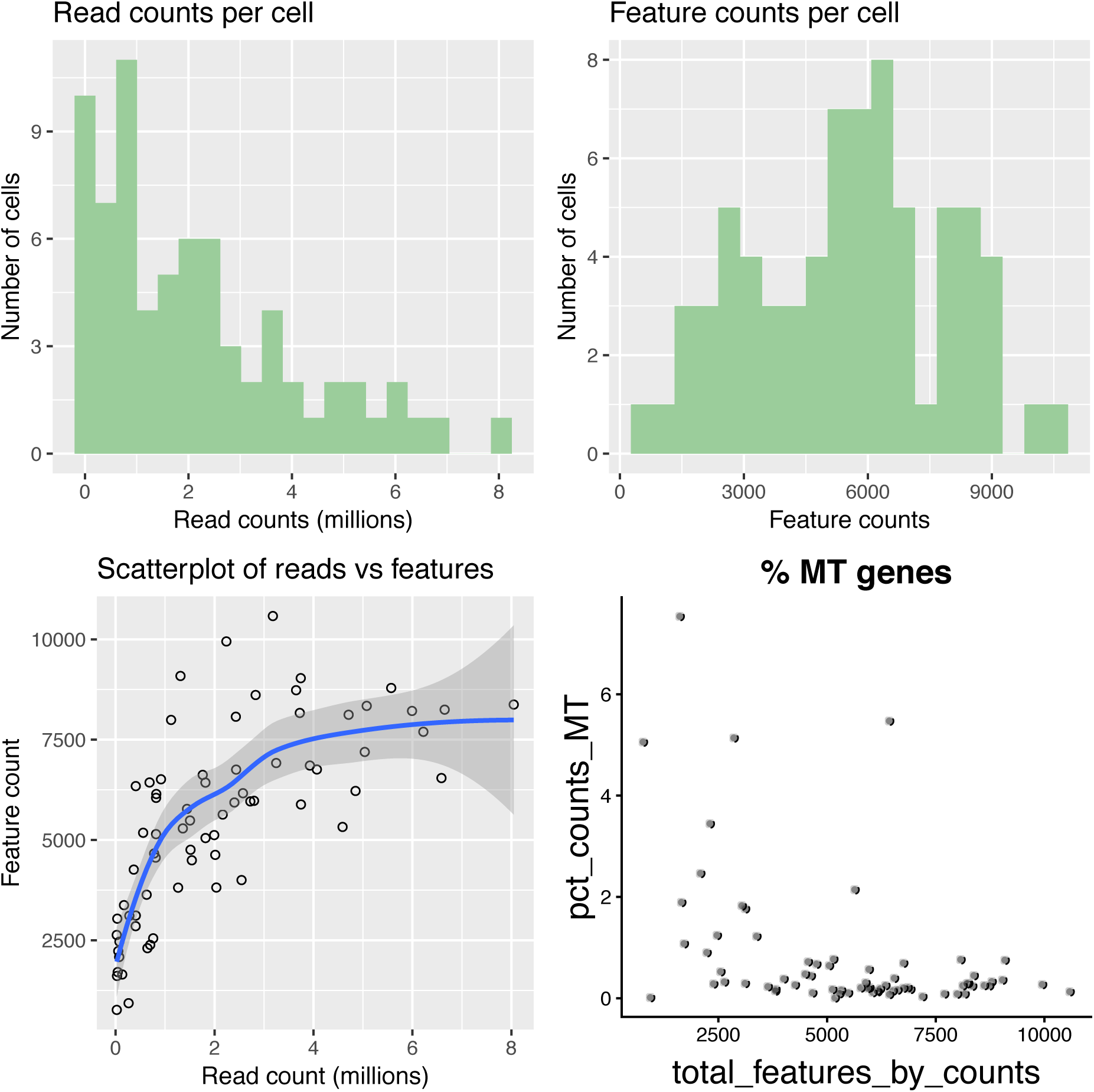
Plotting of data after quality control. Most of the low-quality cells have been removed, leaving cells that have a high number of reads and features (genes in this case) and a low percentage of mitochondrial genes. Note that the tight cluster of cells at the base of the x and y axes in “Scatterplot of reads vs features” has now also disappeared.

## Methods

We present a 5-step workflow in our results above for which we use ‘ready-made’ Scater methods, along with our own bespoke methods.

### Data input

When data is read in, a number of sanity checks take place in order to confirm that the minimum required information has been loaded and then further decisions are made depending on what additional information is loaded (e.g. have ERCC spike-ins been included in the data). This raw data is then used to calculate expression metrics using Scater’s ‘calculateQCMetrics’ method. The output is then saved as a Loom file.

### Plotting tools

There are a number of plotting tools provided to the user. One such tool uses ggplot2 [17] to layout a panel of four plots containing a scatter-plot of mitochondrial gene expression, two histograms depicting read counts and feature counts (genes, transcripts, etc), and finally a scatter-plot of read-counts vs feature counts. An additional tool is provided that uses Scater’s inbuilt ‘plotExprsFreqVsMean’ method, which plots gene expression frequency against mean expression level, in order to examine the effects of technical dropout in the data. Finally, we provide a method to examine batch effect and other confounding factors in our QC’d data. Using Scater’s inbuilt methods, the high-quality data is firstly normalised. Next, a Principle Component Analysis (PCA) is applied to the normalised data, which is then plotted. Points on the plot can then be annotated in relation to any column heading in the experiment annotation file. Categorical data may be given a unique shape or colour, whilst points for continuous data may be given scaled sizes. For example, points on the PCA plot may be coloured by sample, shaped by batch and sized by total features.

### Filtering tool

We provide a filtering tool with 2 alternative methods. In the first one the user manually selects cut-off parameters (usually informed by the plotting tools above), above or below which cells are removed if they do not reach the threshold. The metrics that can be filtered for are: the number of expressed genes, library size (calculated from the number of mapped reads), and percentage of reads mapped to mitochondrial genes. Cells failing these thresholds will be removed. The second method automatically removes cells that are categorised as outliers from PCA. This method works by identifying low-quality cells that have markedly different QC metrics from other cells. Both of the filtering methods are designed to be iterative and the user has the option to re-run filtering from raw data, or refine filtering from a previous filtering step.

## Summary

A large number of tools are available to analyse scRNA-seq data. Many of them require quite advanced computational skills and resources to run. These requirements make it difficult to firstly learn the basics of scRNA-seq analyses and then have the power to perform more complicated downstream analyses on typically large datasets. We have developed tools that make it easy to learn and run typical scRNA-seq quality control steps and analyses. These tools can be used either on the command line in an intuitive, iterative manner, or can be integrated into the Galaxy platform, which will allow further downstream analyses with other tools.

## Availability

All our code and wrappers, complete with installation instructions, tool help and a typical workflow, are available under the MIT open-source license at https://github.com/galaxyproject/tools-iuc/tree/master/tools/scater and on the Galaxy ToolShed at https://toolshed.g2.bx.psu.edu/view/iuc/suite_scater/. The tools can also be freely used at the UseGalaxy.eu public server (https://usegalaxy.eu).

## Acknowledgements

The authors would like to acknowledge the EBI Gene Expression Group (EBI-GEG) for their helpful inspiration, comments and suggestions. The EBI-GEG together with ourselves and other researchers are part of a larger collaborative effort to make Galaxy-based SC analysis more accessible to the community at https://singlecell.usegalaxy.eu/

